# Reprogramming of Cellular Plasticity via ETS and MYC Core-regulatory Circuits During Response to MAPK Inhibition in BRAF-mutant Colorectal Cancer

**DOI:** 10.1101/2025.06.15.659716

**Authors:** Hey Min Lee, Zhao Zheng, Alexey Sorokin, Chi Wut Wong, Stefania Napolitano, Saikat Chowdhury, Preeti Marie Kanikarla, Anand K. Singh, Veena Kochat, Christopher A. Bristow, Sanjana Srinivasan, Michael Peoples, Emre Arslan, Jumanah Yousef Alshenaifi, Oscar E. Villarreal, Van K. Morris, John Paul Shen, Funda Meric-Bernstam, Abhinav K Jain, Natalie Wall Fowlkes, Amanda Anderson, David G Menter, Ajay Kumar Saw, Kunal Rai, Scott Kopetz

**Author notes:** **Corresponding Authors:** Scott Kopetz, MD, PhD, Department of Gastrointestinal Oncology, Unit 0426 The University of Texas MD Anderson Cancer Center 1515 Holcombe Blvd. Houston, TX 77030; Kunal Rai, PhD, Department of Genomic Medicine, Unit 1954, The University of Texas MD Anderson Cancer Center South Campus Research Bldg 4 (4SCR6.1047), 1901 East Road, Houston, TX 77054.

## Abstract

**Background:** Aberrant enhancer dynamics play a critical role in the initiation and progression of colorectal cancer (CRC). BRAF^V600E^-mutated metastatic CRC may be a unique subtype, exhibiting a strong epigenetic phenotype. Interestingly, bromodomain 2, a reader protein for H3K27ac-marked enhancers, was found to be synthetically lethal in CRC with BRAF + EGFR inhibition.

**Design:** We evaluated the effectiveness of targeting aberrant enhancers with bromodomain and extraterminal (BET) + MAPK pathway inhibitors in patient-derived xenograft models of metastatic CRC, followed by comprehensive profiling of transcriptomic and chromatin dynamics upon BET inhibitor combination treatment.

**Results:** The combination of BET and standard MAPK inhibitors has demonstrated improved efficacy against BRAF^V600E^ CRC and selective improvements against RAS-mutant CRC *in vivo*. We showed that BET + MAPK inhibition induced a profound downregulation of the MAPK signaling pathway compared to MAPK inhibition alone. The loss of activation signal on enhancers, as determined by H3K27ac, led to dysregulation of core-regulatory circuitries of CRC, especially loss of the auto-regulatory mechanism of the MAPK downstream E26 transformation–specific transcription factor family. Single nucleus multiome (RNA + ATAC) sequencing further distinguished differential transcriptomic and chromatin dynamics at cell type levels. Profound downregulation of well-differentiated cell types confirmed deep inhibition of MAPK signaling and downstream transcription factors. On the other hand, dedifferentiated cell populations were abundant after MAPK or combination inhibition, suggesting therapy-induced cell state switching and adaptation.

**Conclusion:** We are evaluating BET + BRAF + EGFR inhibition in patients with treatment-refractory BRAF^V600E^ metastatic CRC. ClinicalTrial.gov identifier: NCT06102902.

**What is already known on this topic:** Enhancer aberrations emerge as critical epigenetic features in the progression of colorectal cancer (CRC). However, the dynamics of active enhancer and the therapeutic potential of enhancer blockade, particularly in CRC tumors with BRAF^V600E^ mutation, is not well understood.

**What this study adds:** This study demonstrates improved efficacy of BET inhibitor combination therapies in diverse patient-derived models and reveals epigenetic reprogramming driven cellular plasticity in BRAF^V600E^-mutated CRC.

**How this study might affect research, practice or policy:** Our findings support treatment with BET + BRAF + EGFR inhibitors for patients with BRAFV600E-mutant mCRC [NCT06102902]. This study also highlights the potential of combining epigenetic agents to standard targeted therapy, offering a novel treatment option for this subset of patients.

## Introduction

Colorectal cancer (CRC) is the second leading cause of cancer-related death in the United States, with emerging cases of early-onset CRC ^1,2^. Mutations in the RAS-MAPK pathway are well-known drivers of CRC pathogenesis. *BRAF* mutation occurs in about 5%-10% of patients with metastatic CRC, and most cases are associated with amino acid substitutions of valine to glutamic acid in codon 600 (V600E) ^3^. Additionally, *BRAF* mutations are associated with the consensus molecular subtype1 (CMS1) subtype, which is highly associated with microsatellite instability, the CpG island methylator phenotype, hypermutation, and immune infiltrations ^4–7^. RAS family genes, such as *KRAS* and *NRAS*, are frequently mutated in CRC. *KRAS* mutation occurs in about 40% of cases, predominantly in exon 2, codons 12 (83%) and 13 (14%) ^8,9^. *NRAS* mutations are present in 3%-5% of CRC cases and are mutually exclusive with *KRAS* mutations ^9^. *KRAS* and *NRAS* mutations result in continuous MAPK signaling activation, and patients with these mutations do not benefit from EGFR inhibitor (EGFRi) treatment ^10^.

Many small molecule inhibitors and antibodies have been developed against the key regulators of the MAPK signaling pathway and have been subsequently evaluated in clinical trials and approved as standard therapy regimens for CRC. However, single-drug treatment, such as with the BRAFi, vemurafenib, demonstrated poor prognosis in BRAF^V600E^-mutated CRC, with a < 5% response rate ^11^. Combination therapies have been actively explored to repress intrinsic resistance and emerging acquired resistance to drug treatment, especially MAPK signaling pathway inhibitors. Early studies showed intrinsic resistant feedback activation of EGFR following BRAF inhibition in BRAF^V600E^-mutated CRC ^12^, and the combination of vemurafenib with irinotecan and cetuximab (VIC regimen) has been added to the National Comprehensive Cancer Network (NCCN) guidelines as a therapeutic option for patients with treatment-refractory BRAF^V600E^-mutated CRC ^13,14^. Additionally, the open-label phase III BEACON CRC study showed an improved response rate of 26% for the triplet combination treatment (encorafenib + cetuximab + binimetinib) compared to 20% and 2% for the doublet combination (encorafenib + cetuximab) and control treatments (either cetuximab + irinotecan or cetuximab + FOLFIRI), respectively ^15,16^. Overall survival and progression-free survival durations were most promising for the triplet combination treatment (9.0 and 4.3 months, respectively). Regardless of the promising results of the combination treatment regimens, there is still a strong need to improve therapy response and durability in this subgroup of patients.

Heterogeneity in cancer is a key challenge in precision oncology; tumors must be classified by molecular features for effective treatment, and CMSs are pivotal in the classification of CRC ^4,5^. The emerging significance of epigenetic reprogramming, which involves DNA methylation, histone modifications, and chromatin accessibility, highlights its role in drug sensitivity and resistance mechanisms in cancer ^17–19^. Our previous work introduced enhancer-based chromatin subtypes of CRC, which are correlated with CMS subtypes. Enhancer-based chromatin subtype 1, for instance, is aligned closely with CMS1, showcasing pathways that are associated with immune response^20^. Abnormal histone acetylation levels have been observed in cancer and have been found to have an oncogenic role via transcriptional activation of cancer-related genes ^21^. Depletion of enhancers with bromodomain and extraterminal inhibitors (BETis) in CRC models resulted in varying responses, suggesting the influence of pre-existing enhancer states on treatment outcomes ^20^. All these findings underscore the significance of understanding aberrant epigenomic reprogramming by molecular features or after standard targeted therapies.

The BET family, comprising bromodomain 2 (BRD2), BRD3, BRD4, and BRDT ^22^, recognizes and binds acetylated lysine and acts as an epigenetic reader that regulates transcriptional activation and chromatin remodeling ^22,23^. BET family proteins are implicated in various cancers through the transcriptional regulation of oncogenes such as *MYC* ^24,25^. Small-molecule inhibitors of BRDs have been developed that competitively bind to acetylated lysine recognition pockets and demonstrate a role as anti-cancer agents in various cancers. BETis have demonstrated inhibition of CRC cell proliferation, depletion of c-MYC, and cell cycle arrest in sensitive lines ^26,27^. Additionally, BETis have demonstrated a potential clinical benefit in combination with inhibitors of other oncogenic pathways in many cancer types currently being explored in the clinic.

Overall, continuous efforts are needed to impede the development of resistance to standard therapy and improve response in patients, especially patients harboring the BRAF^V600E^ mutation, which is a unique subgroup that demonstrates a highly epigenetic phenotype. In this study, we evaluated the effectiveness of targeting epigenetic vulnerabilities with BETi and standard MAPK-targeted therapy in patient-derived xenograft models of metastatic CRC. Our data demonstrates promising tumor responses to BETi + BRAFi + EGFRi in BRAF^V600E^ CRC in preclinical patient-derived models and identifies reprogramming of colon cancer cell fate as an underlying mechanism of response and potential resistance.

## Results

### BRAF^V600E^-mutated CRC exhibits a unique enhancer signature

CRC tumors harboring BRAF^V600E^ mutation were characterized by hypermutation, microsatellite instability, and the CpG island methylator phenotype ^6^. We further investigated the potential enhancer dynamics linked to BRAF^V600E^ mutation using publicly available H3K27ac chromatin immunoprecipitation followed by sequencing (ChIP-seq) data obtained from CRC patient tumor samples ^20^. These data comprised four tumor tissues with BRAF^V600E^ mutation and 10 with wild-type RAS/RAF. Certain H3K27ac peaks were differentially bound in the BRAF-mutant or -wild-type samples (318 and 257 sites, respectively; *P* < 0.01) **(Figure 1A)**. Interestingly, significantly elevated H3K27ac active enhancer peaks in BRAF^V600E^ tumors were associated with oncogenic signaling pathways, including the MAPK signaling and PI3K-Akt signaling pathways (adjusted *P* < 0.25) **(Figure 1B)**. This finding indicates that aberrant H3K27ac-marked enhancers may mark oncogenic pathways in BRAF-mutant CRC.

**Figure 1.**
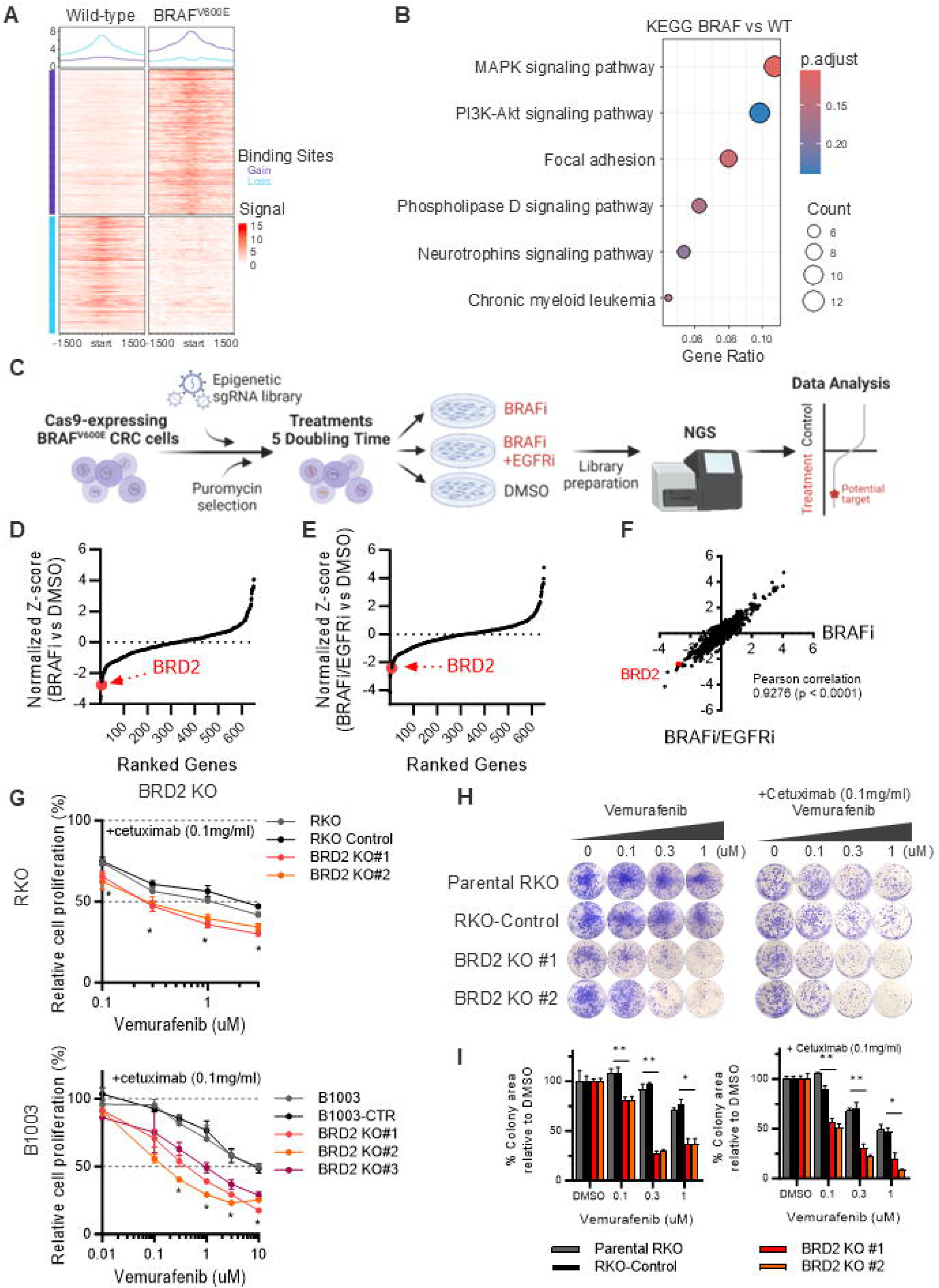
BRD2 deficiency sensitizes BRAF^V600E^ CRC to standard targeted therapy. (**A**) Differential binding of H3K27ac ChIP-seq on BRAF^V600E^-mutated CRC patients compared to wild-type CRC patients (4 and 10 patient tumor samples, respectively) (adjusted *P* < 0.01). (**B**) Pathway analysis using KEGG gene sets on differentially gained H3K27ac peaks in BRAFV600E mutant CRC samples (adjusted *P* < 0.25). (**C**) Schematics of CRISPR library screening design using an epigenetic gene library. (**D**) Genes ranked by the normalized z-score of BRAFi-treated cells relative to DMSO control–treated cells in the RKO cell line; BRD2 as one of the top candidates with synthetic lethality with BRAFi treatment. (**E**) Genes ranked by the normalized z-score of BRAFi + EGFRi–treated RKO cells relative to DMSO control cells; BRD2 as one of the top candidates with synthetic lethality with BRAFi + EGFRi treatment. (**F**) Correlation between normalized z-scores of BRAFi and BRAFi + EGFRi screen in RKO (*P* < 0.0001). (**G**) Relative proliferation of parental, control, BRD2 KO lines in two BRAF^V600E^ CRC cell lines (RKO and B1003) under vemurafenib + cetuximab treatment for 72 hours. \**P* < 0.05 (KO line vs control). (**H**) Colony formation assay of parental, control, and BRD2 KO lines in two BRAF^V600E^ CRC cell lines (RKO and B1003) under vemurafenib + cetuximab treatment for 10 days. (**I**) Relative colony area of parental, control, and BRD2 KO RKO lines upon vemurafenib or vemurafenib + cetuximab treatment for 10 days. \**P* < 0.05, \*\**P* < 0.01.

Further comparisons of H3K27ac enrichment were conducted in CRC tumor tissues harboring BRAF^V600E^ mutation (*n* = 4), KRAS mutation (G12D, G12V, G13D, and L19F; *n* = 7), and wild-type RAS/RAF (*n* = 10). The higher number of H3K27ac peaks were differentially enriched in either BRAF-mutant or wild-type RAS/RAF CRC tumors compared to KRAS-mutant tumors **(Supplemental Figure 1, A and B)**. KRAS-mutant tumors demonstrated more H3K27ac enrichment around differentially bound sites of wild-type RAS/RAF tumors than BRAF-mutant tumors **(Supplemental Figure 1C)**. Overall, this result indicates unique H3K27ac-marked active enhancer profiles among BRAF^V600E^ CRC tumors.

### BRD2 deficiency improves response to BRAF/EGFR inhibition

On the basis of the robust correlation with DNA hypermethylation and aberrant enhancer aberrations linked to BRAF mutation, indicating that BRAF^V600E^-mutant CRC represents a highly epigenetically modulated subtype, we hypothesized that targeting the key epigenomic susceptibility would enhance the sensitivity of BRAF^V600E^-mutated tumors to standard targeted therapies. We conducted CRISPR-Cas9 drug modifier screening in the Cas9-expressing BRAF^V600E^-mutated CRC cell line RKO with a library of 519 genes that are associated with epigenomic regulation ^28,29^ to unbiasedly identify candidate genes that have a synthetically lethal relationship with vemurafenib (BRAFi) or vemurafenib + cetuximab (BRAFi + anti-EGFR antibody) treatment **(Figure 1C)**. Intriguingly, BRD2 was identified as one of the top hits that are synthetically lethal under both vemurafenib and vemurafenib + cetuximab combination treatment **(Figure 1, D and E)**. Intriguingly, the synthetic lethality of genes was highly similar under vemurafenib or vemurafenib + cetuximab treatment **(Figure 1F)**.

We further validated whether genetic knock out (KO) of BRD2 could sensitize BRAF-mutant cell lines to BRAFi + EGFRi treatment. BRD2 was knocked out in two BRAF^V600E^-mutated CRC cell lines, RKO and B1003, and treated with either vemurafenib or vemurafenib + cetuximab. A cell viability assay confirmed that BRD2 deficiency significantly sensitized cancer cells to both monotherapy and combination therapy treatment **(Figure 1G)**. BRD2 KO clones also became deficient in colony formation compared to parental and negative control lines **(Figure 1, H and I)**. These data suggest that BRD2 plays an oncogenic role in BRAF^V600E^ CRC and that its depletion is synthetically lethal with targeted therapy.

### Combining BETis improves the efficacy of standard targeted therapies in CRC in vivo

BRD2 activity through binding to histone acetylated lysine residue can be blocked by BETi treatment, and there are multiple BETis under development that demonstrate anti-cancer efficacy. Thus, we assessed whether combining BETi with BRAFi + EGFRi can improve efficacy using BRAF^V600E^-mutated CRC patient-derived xenograft (PDX) models. Excitingly, significant tumor regression was observed with triplet combination treatment (ZEN-3694 [BETi] + encorafenib [BRAFi] + cetuximab [EGFRi]) in all three PDX models compared to either ZEN-3694 or encorafenib + cetuximab treatment alone **(Figure 2A)**; mice tolerated the triplet combination without noticeable body weight changes **(Figure 2B)**. Although responses in one of the models (C0999) under triplet combination treatment were heavily driven by encorafenib + cetuximab, tumor regression was only observed upon triplet combination treatment in the other two models (C5002 and B8120); stable tumor growth was observed with individual drug treatment.

**Figure 2.**
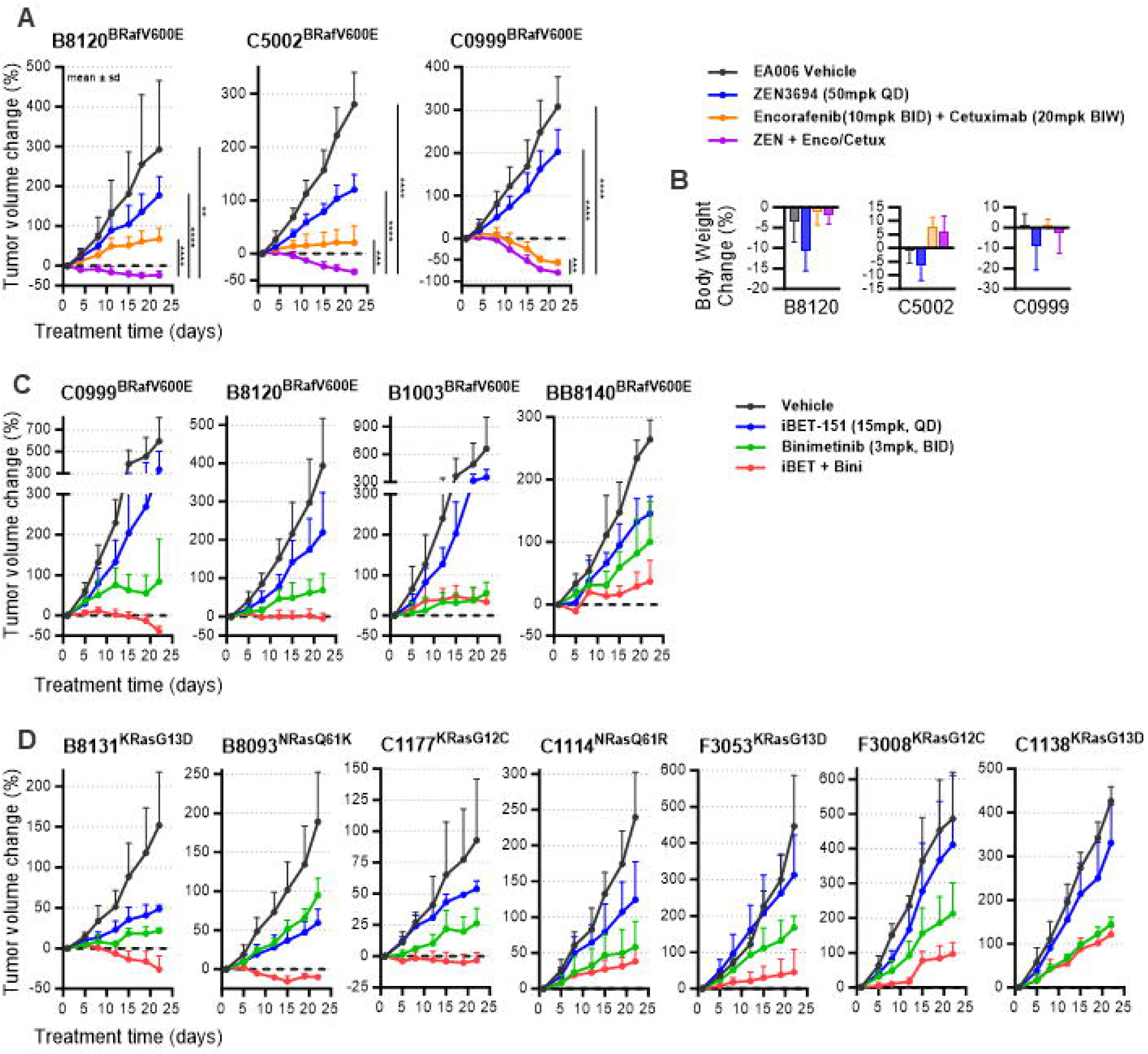
BETi combination with standard targeted agents for MAPK blockade improves efficacy in BRAF^V600E^ CRC PDX models. (**A**) Tumor volume changes upon treatment with vehicle control, ZEN-3694 (BETi), encorafenib (BRAFi) + cetuximab (EGFRi), and triplet combination treatment relative to baseline in three BRAF^V600E^-mutated CRC PDX models (mean, *n* = 7-8 tumors per condition). (**B**) Body weight changes upon treatment in each PDX model (mean, *n* = 7-8 tumors per condition). (**C**) Tumor volume changes upon vehicle control, iBET-151 (BETi), binimetinib (MEKi), and iBET-151 + binimetinib treatment across BRAF^V600E^ CRC PDX models and (**D**) RAS-mutant CRC PDX models (mean, *n* = 5 tumors per condition).

On the basis of the promising responses observed in BRAF^V600E^-mutant CRC PDXs, we extended the assessment of the efficacy of BETi combinations in RAS-mutant CRC PDX models. Despite the limited treatment options available for RAS-mutant CRC, efforts to target these tumors have been primarily aimed at the RAS-MAPK cascade. Thus, we switched the backbone MAPK inhibition to MEKi (binimetinib) to assess its efficacy in BRAF-mutated, RAS-mutated and -wild-type CRC models.

iBET-151 (pan-BETi) plus binimetinib resulted in tumor regression in two BRAF^V600E^ CRC and three RAS-mutated CRC PDX models **(Figure 2C and D)**. We further evaluated the efficacy of this treatment in KRAS-mutated PDAC PDX models. Heterogenous responses were observed **(Supplemental Figure 2)**, which indicates the synergy of this treatment in other cancer types as well.

### Deconvoluting BETi + MEKi treatment

Our previous work showed that H3K27ac occupancy decreased in a larger number of enhancers across the genome in CRC PDXs after BETi or BETi + MEKi treatment ^20(pp938–949)^. Nonetheless, the role of enhancer enrichment and its underlying effects on certain signaling pathways requires further understanding in CRC. In that study, we used H3K27ac binding data to de-convolute the activity of PDX models (B8131, F3008, and C1138) treated with vehicle, iBET-151, trametinib (MEKi), or iBET-151 + trametinib ^20(pp938–949)^.

To understand how enhancer depletion by BET inhibition influences tumor biology, we isolated the unique H3K27ac peaks that were present in both control and MEKi-treated tumor samples that are lost upon BETi or BETi + MEKi treatment, respectively **(Figure 3A)**. We identified 92 genes that commonly lost histone acetylation after BET inhibition across PDX models. Interestingly, these genes were significantly enriched in RAS-MAPK signaling– associated gene sets, such as prolactin activation of MAPK signaling, KRAS signaling UP, and the Ras signaling pathway **(Figure 3A)**. Previous studies demonstrated the accumulation of enhancers around MAPK signaling–related genes, such as *EGFR*, *ERBB2*, *NRAS*, or *KRAS* in CRC tumors compared to normal tissues ^20,30^. Our findings reveal reductions in enhancer activity around genes that are implicated in MAPK signaling pathways following BETi treatment, demonstrating the therapeutic promise of enhancer-targeting in CRC.

**Figure 3.**
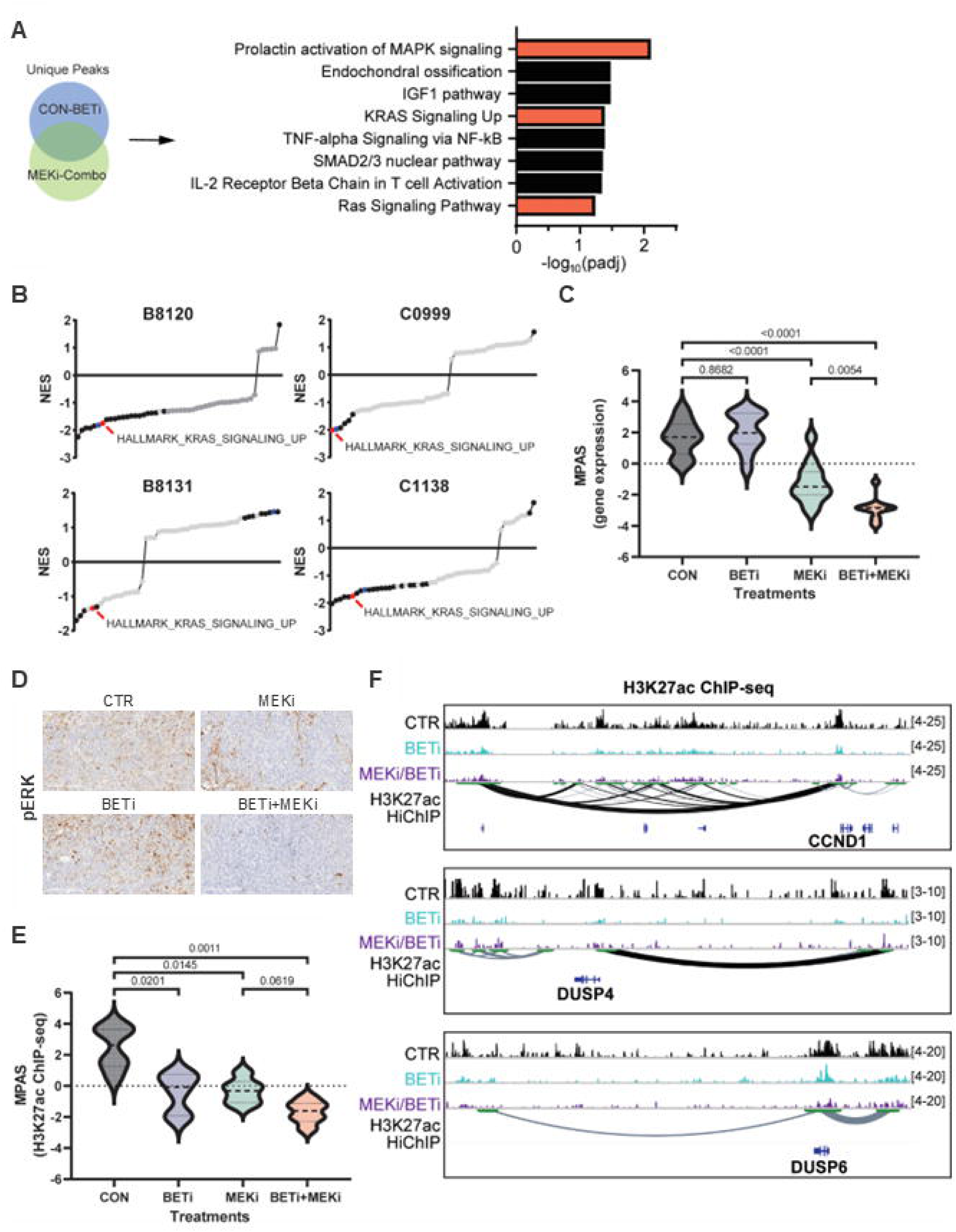
RAS-MAPK signaling pathway inhibition via BETi + MEKi at genetic and epigenetic levels. (**A**) H3K27Ac ChIP-seq peaks that were unique to CTR relative to BETi (CTR-BETi), as well as to MEKi relative to BETi/MEKi (MEKi-Combo), were annotated with genes and then compared between PDX models (B8131, C1138, and F3008). Gene set enrichment analysis of the common genes with enhancer loss after treatment. RAS-MAPK-associated gene sets are highlighted in red. BioCarta, BioPlanet, and Hallmark gene sets were used. (**B**) Gene set enrichment analysis performed with Hallmark gene sets between BETi + MEKi compared to MEKi using RNA-seq data from four PDX models (B8120, C0999, B8131, and C1138). Gene sets were ranked by normalized enrichment score (NES), and significant gene sets (*P* < 0.01) are highlighted in black. The ‘Kras signaling UP’ gene set is highlighted with red. The ‘TNFα signaling via NFκB’ gene set is highlighted in blue. (**C**) MAPK pathway activation score (MPAS), calculated from gene expression under control, BETi, MEKi, or BETi + MEKi treatment in PDXs. (**D**) An Immunohistochemistry (IHC) staining (20x) identifying the pERK in C0999 PDX tissue among four treatments. (**E**) MPAS, calculated from H3K27Ac ChIP-seq data, compared among control, BETi-, MEKi-, or BETi + MEKi–treated PDX tumors. (**F**) IGV snapshots of key MAPK signaling pathway genes, such as *CCND1*, *DUSP4*, and *DUSP6*, from H3K27Ac ChIP-seq from B8131 PDX tumors treated with control, BETi, MEKi, or BETi + MEKi. HCT116 H3K27Ac HiChIP-seq revealed enhancer-promoter loops that are associated with these genes.

### Deep inhibition of MAPK signaling with BETi combination treatment

To better understand the adaptive changes to treatment, we further examined the transcriptomic changes in signaling pathways after short-term (7 days) treatment. Pathway modulation by different therapies in PDX models confirmed the downregulation of KRAS signaling under MEKi relative to control, and surprisingly, under BETi + MEKi relative to MEKi alone **(Figure 3B)**. Significant downregulation of the transcriptional MAPK signaling activation score (MPAS) ^31^ with BETi + MEKi compared to MEKi alone indicated deep inhibition of transcriptional MAPK pathway activation, while no significant difference was observed with BETi alone **(Figure 3C)**. BETi + MEKi demonstrated enhanced suppression of phospho-ERK (pERK) expression, a marker of MAPK signaling activity, in PDXs compared to MEKi alone **(Figure 3D)**. Limited inhibition of pERK indicates sustained activation of MAPK signaling and suggests potential adaptive or acquired resistance to standard MAPK inhibition in CRC. The findings in this study suggest that combined inhibition of BET and MEK induces a more profound blockade of the RAS-MAPK signaling pathway, highlighting the clinical advantages of this approach.

The depletion of H3K27ac peaks around MAPK pathway target genes ^31^ was confirmed following treatment with BETi and MEKi, alone and combined **(Figure 3E)**. Relative downregulation was observed with BETi + MEKi compared to MEKi alone, but it had limited significance. Nonetheless, consistent downregulation, along with a notable loss of H3K27ac, was found around MAPK pathway genes under combined treatment in PDX models, such as *CCND1*, *DUSP4*, and *DUSP6* **(Figure 3F)**. We further highlighted the enhancer-promoter interactions around these MAPK target genes using HCT116 Hi-ChIP data ^20^, validating that the loss of these enhancers can affect the transcription of these genes. Depletion of acetylation peaks was observed around the transcription site as well as looped enhancer sites **(Figure 3F)**. Taken together, BETi + MEKi resulted in an enhanced blockade of the MAPK signaling pathway compared to MEKi alone, as evidenced by the extent of expression changes following the loss of enhancer peaks around key MAPK pathway genes.

### BETi-mediated repression of c-Myc in CRC

On the basis of the depletion of H3K27ac, which is defined as an active enhancer mark, upon BETi or combination treatment in PDXs, we hypothesized that BET inhibition would modulate the activity of the transcription factors (TFs) associated with the MAPK signaling pathway ^32,33^. The results of previous studies have suggested that MYC is a target of BET inhibition across different cancers ^24,25^. The oncogenic activation of MYC has been recognized in cancer cell proliferation and survival ^34^, and MYC amplification is commonly observed in human cancers ^35^; however, therapeutically, targeting MYC has been challenging. Our data confirmed significant downregulation of MYC expression by BETi in three of four PDX models **(Figure 4A)**. Intriguingly, a decrease in MYC expression by BETi synergized with MEKi treatment in all models (synergy index (SI) > 0 ^36^).

**Figure 4.**
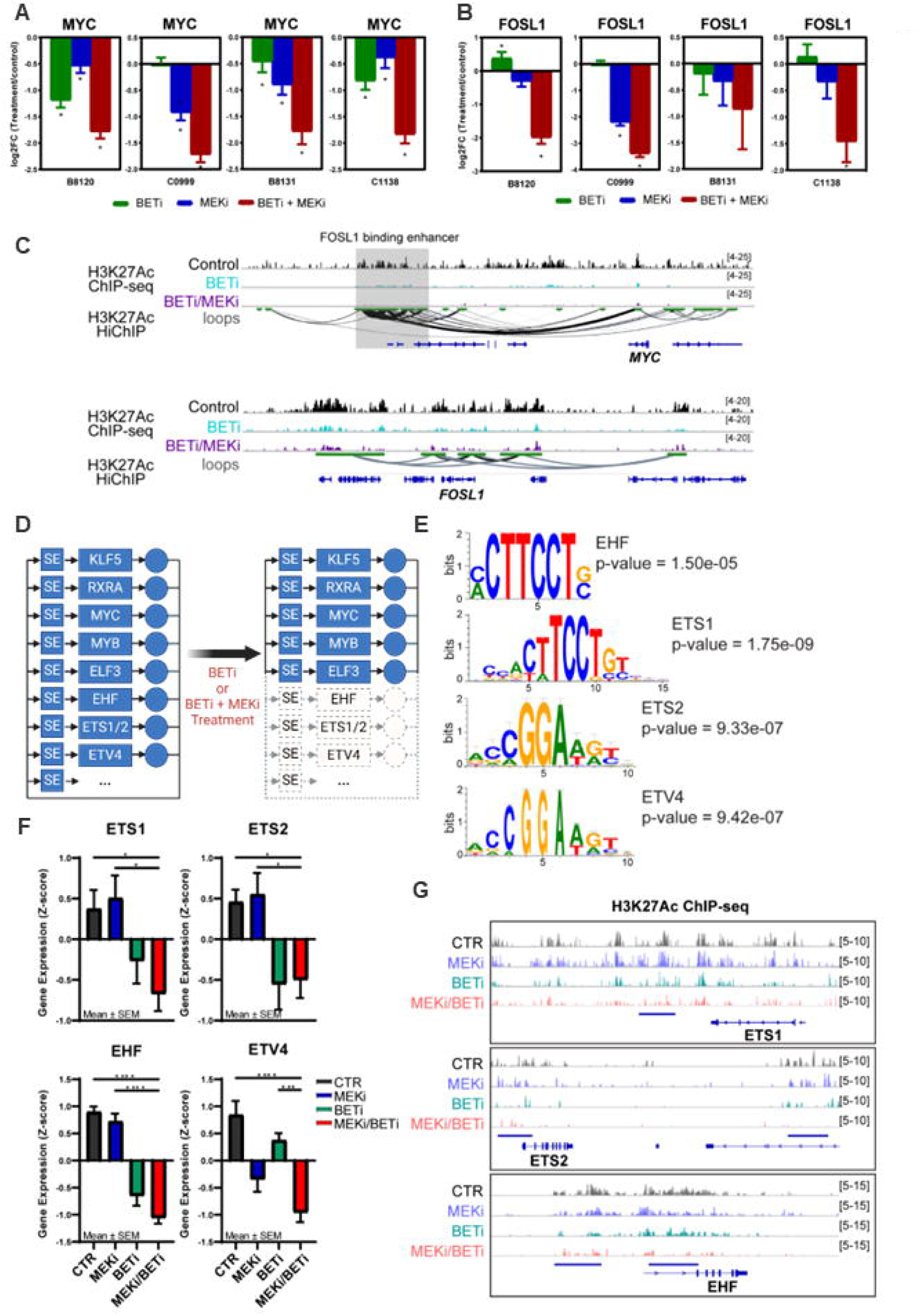
BETi or BETi + MEKi reprograms enhancer-dependent auto-regulated TFs downstream of the MAPK signaling pathway. (**A**) Gene expression changes of MYC upon treatment relative to control in each PDX model. \**P* < 0.05. (**B**) Gene expression changes of FOSL1 after treatment relative to control in each PDX model. \**P* < 0.05. (**C**) IGV snapshot of H3K27ac ChIP-seq from the B8131 PDX treated with BETi, MEKi, and BETi + MEKi. Enhancer-promoter loops from HCT116 H3K27ac HiChIP-seq highlighting enhancer regions associated with MYC and FOSL1. MYC enhancer regions that overlapped with the FOSL1 binding site are highlighted in gray. (**D**) Auto-regulatory TFs commonly found across PDX models. The ETS family is highlighted with dotted lines and was inhibited under BETi or BETi + MEKi treatment. (**E**) Motif analysis of ETS family genes after control or MEKi treatment in F3008 PDX. (**F**) Expression of ETS family under control, BETi, MEKi, or BETi/MEKi treatment. \**P* < 0.05, \*\**P* < 0.01, \*\*\**P* < 0.001, \*\*\*\**P* < 0.0001. (**G**) IGV snapshot of H3K27Ac enhancer enrichment under control, BETi, MEKi, and BETi + MEKi treatment in the F3008 PDX model and associated super enhancer sites highlighted with blue lines.

The results of previous studies suggested that certain TFs share binding sites around super-enhancers (SEs) upstream of the MYC transcription site in CRC ^26,37^. FOSL1, which is an ERK target TF, was also found to bind to the MYC SE site ^37^. We hypothesized that enhancer depletion upon BET inhibition around the MYC transcription site would dysregulate the binding of ERK target TFs, such as FOSL1, and cause synergistic MYC repression. We observed downregulated FOSL1 expression by either MEKi or BETi + MEKi treatment **(Figure 4B)**. Additionally, our data showed a profound depletion of H3K27ac peaks around MYC and previously reported shared genomic regions between MYC SEs and FOSL1 binding sites ^26,37^ after BETi or BETi + MEKi treatment **(Figure 4C)**. We further highlighted enhancer-promoter interactions around MYC using HCT116 Hi-ChIP data, validating that the loss of these enhancers affects MYC transcription. Overall, this result indicates that BETi-induced disruption of enhancer regions, where MAPK pathway–related TFs bind, leads to the downregulation of both MYC and FOSL1 in combination treatment.

### BET inhibition reprogrammed the core-regulatory TFs of CRCs

Depletion of H3K27ac peaks was observed at the global level and around key TFs, so we further assessed modulation in SE–regulated TFs after BETi treatment. Studies suggested that SE-driven core TFs can form interconnected auto-regulatory loops that collectively control gene expression and dysregulate disease-associated transcription ^38,39^. We hypothesized that enhancer blockade with BET inhibition could modify the regulatory mechanism of enhancer-driven TFs.

We identified the auto-regulated core TFs using H3K27ac ChIP-seq data from CRC PDX models. We first classified the list of top core TFs associated with control and MEKi-treated PDX tumors by filtering out TFs that were autoregulated in at least 1/3 of control or MEKi-treated tumor samples (≥ 4 of 12 total samples) **(Figure 4D)**. We further narrowed down the list by core-regulatory TFs in CRC cell lines (HT29, HCT116, SW480, and COLO205) using publicly available H3K27ac ChIP-seq ^39^ **(Supplemental Figure 3A)**. Consequently, several TFs, including KLF5, retinoid X receptor (RXR)A, MYC, MYB, ELF3, and EHF, were identified as core TFs that are critical for the maintenance of CRC tumorigenesis or progression. KLF5, MYC, ELF3, EHF, and other E26 transformation–specific (ETS) TFs, such as ETV4 and ETS2, were previously identified as colon cancer–driving master TF candidates that are regulators of tumor cell states, and RXRA and MYB were identified as master TFs in other cancers, including breast and lung cancer ^40^. A recent study also demonstrated that KLF5 and co-factors (BRD4, MED1, and RAD21) construct the core-regulatory circuitry in the 3D genome structure in CRC ^41^.

To support our hypothesis, we classified TFs that were no longer self-regulated after BETi and BETi + MEKi treatment in at least two PDX models. We found ETS homologous factor (EHF) lost SE–driven autoregulatory features after both BETi and BETi + MEKi treatment. We expanded our investigation to other ETS TFs and found that ETS1/2 and ETV4 also lost autoregulation with BETi in at least one PDX model **(Figure 4D)**. The self-regulatory pattern of ELF3, on the other hand, was maintained in certain PDX models. Motifs found on SE-associated TFs, including the ETS TF family, are highlighted in **Figure 4E** and **Supplemental Figure 3B**. ERK-phosphorylated ETS TFs regulate RAS-MAPK target gene activation and play a role in the proliferation, invasion, and migration of cancer cells.

EHF regulates the human epidermal growth factor receptor family, modulating AKT or MAPK signaling in other cancers ^42–45^ and regulates transforming growth factor β signaling in CRC ^46^. The ETS1/2-containing TF-enhancer complex was found to be regulated by AKT- or MAPK-induced CBP/p300 phosphorylation ^47,48^ or co-localization of the complex with BRD2/4 ^49,50^, supporting the loss of enhancer-driven auto-regulation of ETS1/2 upon BRD inhibition. Additionally, activator protein 1 binding sequences are frequently located near ETS binding sites around promoter regions of MAPK target genes, such as CCND1, which may activate RAS-MAPK signaling in the absence of direct pathway activation ^51–53^. Overall, the ETS family could be a therapeutic target in CRC, and loss of ETS self-regulation upon BETi treatment could additively block the RAS-MAPK pathway.

We further validated the decreased expression of EHF, ETS1/2, and ETV4 under BETi or BETi + MEKi treatment in PDX models **(Figure 4F)**. BETi or BETi + MEKi treatment induced loss of H3K27ac signals on proximal and distal enhancers associated with ETS family genes, while no significant changes were observed with MEKi treatment **(Figure 4G)**. Taken together, our data suggest that ETS gene activation is modulated through a loss of enhancer signals by BET inhibition, leading to additive downregulation of RAS-MAPK signaling in combination with MEK inhibition.

### Differential tumor cell types’ responses upon therapeutic stress

Deep inhibition of the MAPK signaling pathway was also validated in BRAF^V600E^ CRC cell lines after BETi + BRAFi + EGFRi treatment, as assessed by downregulation of pERK expression andDUSP6 gene expression **(Figure 5A and B)**. The results of recent studies suggest that tumor heterogeneity influences responses or resistance to therapy and that diverse clonal fate and phenotype switching occur upon therapeutic stress ^54,55^; however, how heterogeneous cell types respond to targeted therapy, particularly under BET inhibition, is not yet explained in CRC.

**Figure 5.**
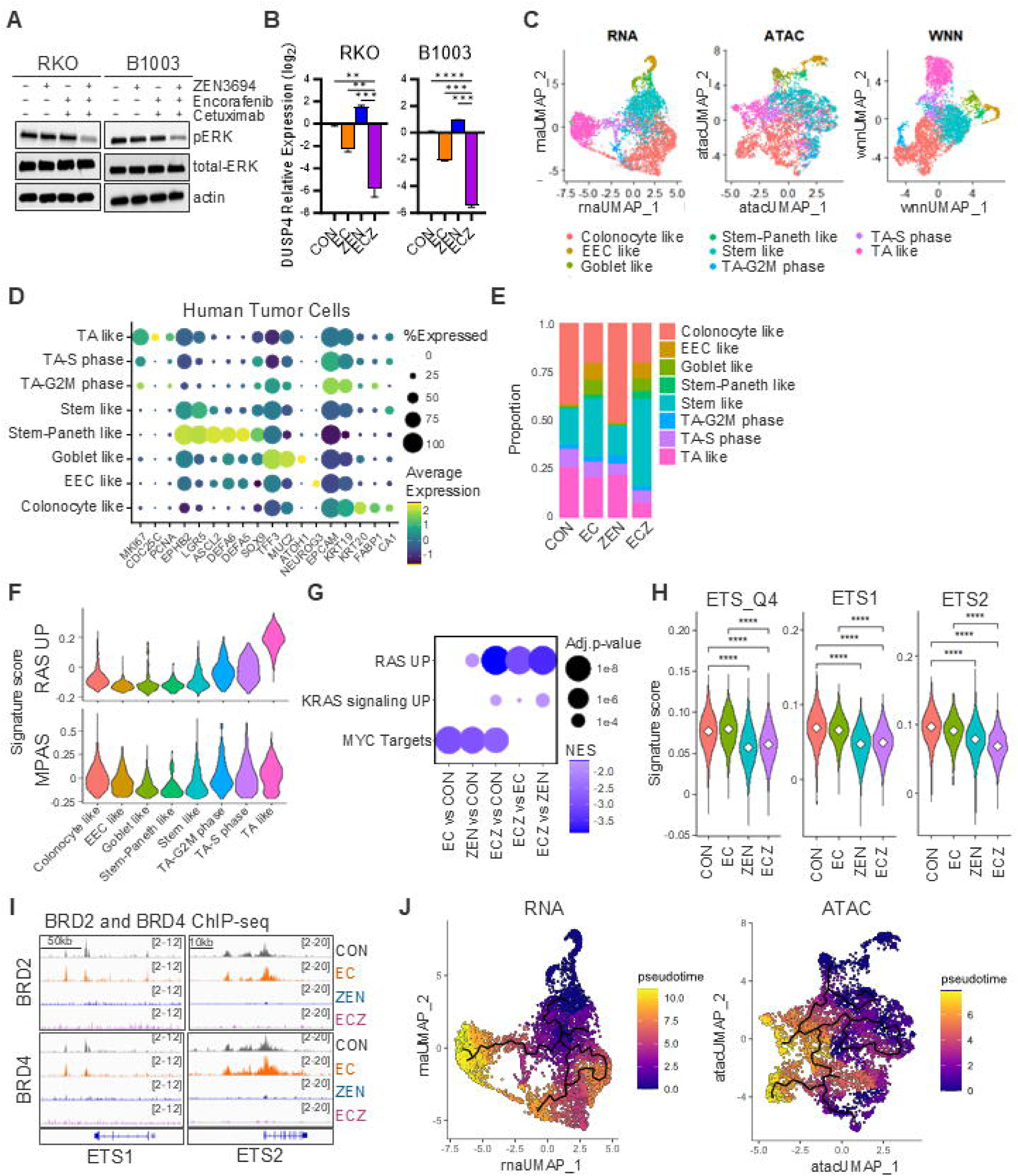
snMultiome sequencing revealed differential responses to therapy by tumor cell type. (**A**) A western blot analysis demonstrated expression of phosphor ERK, total ERK, and actin in BRAF^V600E^ CRC cell lines (RKO and B1003) treated with ZEN-3694 (3μM), encorafenib (0.5μM), or cetuximab (0.01mg/ml) for 10 hours. (**B**) Relative expression of DUSP6 in RKO and B1003 cell lines treated with DMSO (CON), encorafenib + cetuximab (EC), ZEN-3694 (ZEN), or encorafenib + cetuximab + ZEN-3694 (ECZ) for 72 hours with GAPDH as reference gene. (**C**) UMAP projection of human tumor cells, annotated on the basis of the expression of key epithelial cell marker genes in RNA, ATAC, and merged WNN data from snMultiome (RNA+ATAC) sequencing from C5002 PDX tumor tissues. (**D**) Dot plot representing the expression of key marker genes across cell types. Percentage (%) of expression in cells is represented by dot size and colored on the basis of the average expression of each gene in the C5002 PDX tumor snMultiome data. (**E**) Abundance of each types cell population by treatment groups including control (CON), encorafenib + cetuximab (EC), zen3694 (ZEN), and encorafenib + cetuximab +zen3694 (ECZ). (**F**) RAS UP signature and MAPK activation scores (MPAS) across cell types in C5002 PDX tumor snMultiome data. (**G**) Gene set enrichment analysis (GSEA) of ‘RAS signature UP’, hallmark ‘KRAS signaling UP’, and hallmark ‘MYC Targets’, conducted by comparing treatments groups, including vehicle (CON), encorafenib + cetuximab (EC), ZEN-3694 (ZEN), and encorafenib + cetuximab + ZEN-3694 (ECZ), in C5002 PDX tumor snMultiome data. Dot size represents adjusted *P* values, colored by the normalized enrichment score (NES). (**H**) ETS target gene signature scores were calculated for ETS_Q4, ETS1, and ETS2 target gene sets (GSEA C3:TFT) by treatment groups in the C5002 PDX tumor snMultiome data. \*\*\*\**P* < 0.0001. (**I**) IGV snapshot of BRD2 or BRD4 peaks around ETS1 and ETS2 transcription sites in C5002 PDX tumor samples. (**J**) UMAP projection of trajectories calculated from Monocle3 in RNA and ATAC-seq data. Using stem-paneth like cell cluster as a starting node, each cluster were colored by calculated pseudotime representation of differentiation.

Tumor tissues from one BRAF^V600E^ PDX model (C5002, **Figure 2A**), which achieved regression only upon triplet combination treatment, were profiled using single-nucleus Multiome (snRNA + ATAC) sequencing to determine how tumor cells are differentially modulated under ZEN-3694 + encorafenib + cetuximab treatment compared to ZEN-3694 or encorafenib + cetuximab treatment alone. Tumors were collected after 10 days of treatment, when they had started to regress under triplet combination treatment, to ensure a sufficient therapeutic effect and sufficient tumor cells for sequencing. First, cells originating from human tumor epithelial cells were isolated from a mixture of mouse cells, and we assessed treatment effects using only human tumor cells. All clusters of cells were identified as epithelial cells using SingleR ^56^. We further differentiated the clusters by their similarities to intestinal cell subtypes on the basis of the expression of key genes **(Figure 5C and D)**. Interestingly, therapeutic stress following encorafenib + cetuximab or the triplet combination treatment successfully depleted the proliferative and well-differentiated (TA- or colonocyte-like) cell populations **(Figure 5E)**.

Activation of the RAS-MAPK signaling pathway was compared between cell types using robust, clinically relevant gene signatures (RAS UP) ^57^ and Hallmark RAS signaling UP; higher activity was demonstrated in proliferative TA- or colonocyte-like cell populations, which were most abundant in control and BETi-treated cells **(Figure 5F, Supplemental Figure 4)**. Enrichment of the RAS-MAPK signaling pathway was significantly downregulated with BETi + BRAFi + EGFRi treatment compared to BETi or BRAFi + EGFRi treatment **(Figure 5G)**. The enrichment of ETS TF target gene sets was significantly decreased upon BETi or BETi + BRAFi + EGFRi treatment **(Figure 5H)**.

A significant decrease in MYC target enrichment was observed upon treatment **(Figure 5G)**. MYC and FOSL1 were mostly expressed around proliferative TA-like cells, and associated motifs (MA0147.3 and MA0477.2) were found to be enriched in TA- or colonocyte-like cell populations which are depleted upon treatment **(Supplemental Figure 5A and B)**. Downregulation of MYC and FOSL1 expression upon BETi + BRAFi + EGFRi treatment was confirmed **(Supplemental Figure 5C)**. A profound depletion of BRD2 and BRD4 binding upon BETi treatment was also confirmed in PDX tumor tissues **(Supplemental Figure 5D and E)**, and found to be significantly associated with MYC targets, supporting MYC downregulation by BETi treatment in our PDX model **(Supplemental Figure 5F and G)**. Additionally, loss of BRD2 and BRD4 binding was confirmed around ETS TFs **(Figure 5I)**, indicating successful BRD blockade with BETi and a potential role for BRDs in regulating ETS TFs. Thus, both BRD2 and BRD4 blockade with BETi treatment is critical in regulating RAS signaling as well as MYC target genes.

Differential responses to therapy were observed in different cell populations. Predicted trajectories identified by monocle3 further distinguished cell differentiation states by cell types in both snRNA-seq and snATAC-seq **(Figure 5J)**. Interestingly, less-differentiated cell populations that highly express intestinal stem maker genes were abundant upon targeted therapy **(Figure 5E)**, indicating treatment-induced cell state switching and potential tumor adaptation and resistance. This result provided hypothesis for translational studies in the clinical trial and gives us leads to understand treatment resistance in patients.

### BET inhibition may help overcome acquired resistance to standard targeted inhibition of MAPK signaling

We confirmed improved efficacy in treatment-naïve PDX models upon combined BET inhibition and further assessed it in resistant PDX models. We established acquired-resistance PDX models through continuous treatment with BRAFi + EGFRi until tumor growth was observed. First, the combination of BRAFi + EGFRi + BETi significantly reduced the proliferation compared to either BETi or BRAFi + EGFRi in PDX-derived organoid (PDXO) models, which had acquired resistance to encorafenib + cetuximab **(Figure 6A and B)**. Additionally, the efficacy of BETi with two different MAPKi regimens, BRAFi + EGFRi or MEKi, was assessed in another resistant PDX model that contained an acquired KRAS^G12D^ mutation and resistance to vemurafenib + cetuximab, mimicking what is observed in patients. Considering the aggressiveness of the resistance model, the BETi combination significantly stabilized tumor growth compared to individual drug treatment **(Figure 6C and D)**. This finding suggests a potential role for BETi combination therapy in overcoming resistance to standard therapy in BRAF^V600E^-mutant CRC, which could be a promising therapeutic regimen for patients with residual disease.

**Figure 6.**
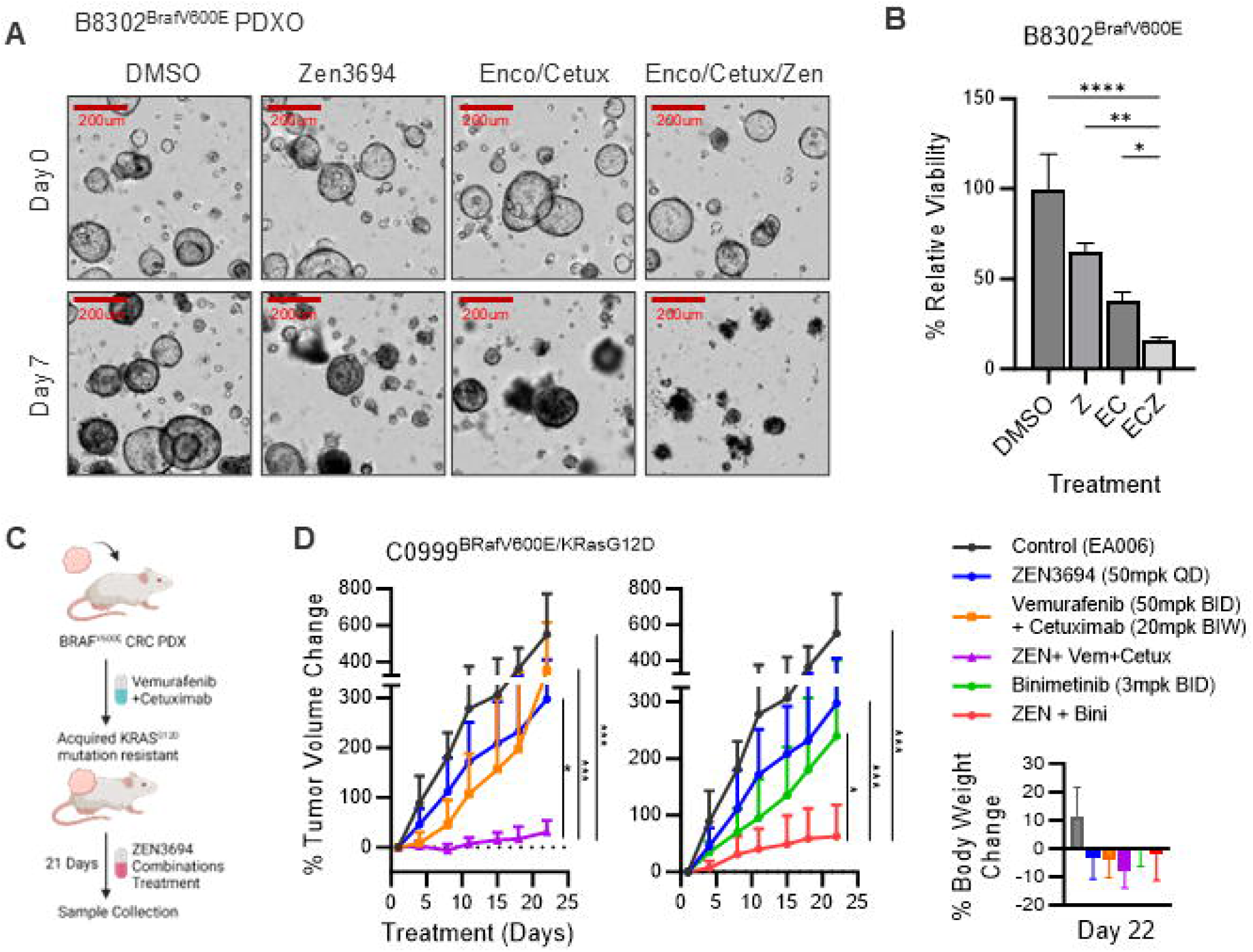
Combined BETi treatment sensitized BRAF-mutant PDX models that had acquired resistance to BRAF or EGFR inhibition. (**A**) A resistant PDXO (B8302) model at the start (day 0) and endpoint of treatment (day 7) with DMSO, ZEN-3694 (3 µM), encorafenib + cetuximab (Enco/Cetux; 0.05 µM/0.1 mg/ml), or encorafenib + cetuximab + ZEN-3694. (**B**) Viability of PDXO (B8302) under treatment with ZEN-3694, encorafenib + cetuximab, or encorafenib + cetuximab + ZEN-3694 relative to DMSO control, as measured by luminescence. \**P* < 0.05, \*\**P* < 0.01, \*\*\**P* < 0.001, \*\*\*\**P* < 0.0001. (**C**) An illustration of the experimental design of an acquired resistance PDX model. (**D**) Percentage of the tumor volume change of C0999, vemurafenib + cetuximab–resistant PDX, up to 21 days after treatment with control, ZEN-3694, vemurafenib + cetuximab, the triplet combination, binimetinib, or the doublet combination treatment. Percentage of body weight changes as a measure of toxicity under different treatment conditions at the endpoint of treatment. \**P* < 0.05, \*\**P* < 0.01, \*\*\**P* < 0.001.

## Discussion

Aberrant chromatin dynamics plays a critical role in the initiation and progression of CRC. In this study, we designed a combined treatment regimen that is specific to the subgroup of metastatic CRC patients bearing BRAF^V600E^ mutation, which exhibits an epigenetic phenotype.

This study identified a synthetically lethal relationship between BRD2 and BRAF or BRAF or EGFR inhibition in BRAF^V600E^-mutated CRC. Combined inhibition of BRD and the MAPK signaling pathway showed successful tumor regression in treatment-naïve and acquired-resistance BRAF^V600E^ CRC PDXs, as well as RAS-mutant CRC PDXs. The findings from this study demonstrated deep inhibition of the MAPK signaling pathway with BET plus MAPK blockade compared to MAPK pathway blockade alone at both epigenetic and transcriptomic levels. Enhancer depletion by BET inhibition dysregulated enhancer-dependent autoregulation of downstream TFs of the MAPK pathway, particularly the ETS family. Additionally, the single nucleus multiome sequencing results suggest successful depletion of proliferative and differentiated tumor cells that highly express MAPK signaling or MYC targets with triplet combination treatment; however, we observed enrichment of stem-like tumor cells upon BRAF + EGFR or BET + BRAF + EGFR inhibition, which suggests treatment-resistant plasticity. Overall, the results of this study indicate that the novel combination of BETi + BRAFi + EGFRi treatment will improve responses in BRAF^V600E^ CRC patients.

The results of this study emphasize that BETi induces epigenetic reprogramming of MAPK downstream TFs, leading to more profound inhibition of the MAPK signaling pathway. It has been suggested that nearly complete inhibition of phospho-ERK is necessary for optimal clinical effectiveness in MAPK pathway–targeted therapy ^58^. However, achieving such complete inhibition of ERK activity in CRC is challenging ^59^, even with combinations of different MAPK pathway inhibitors showing some clinical activity. This study suggested that combined BET inhibition is a promising therapeutic approach to enhancing MAPK signaling blockade and reducing ERK activity, improving clinical efficacy. An extended investigation of BETi combinations in RAS mutant solid tumors, including CRC, is worth further consideration; one such study is currently enrolling patients (NCT05111561).

CRC tumors with BRAF^V600E^ mutation exhibit a relatively high mutation burden and prevalence in CMS1, which is characterized as an immune-active subtype ^4^. A study suggest that MAPK pathway blockade by BRAF or EGFR inhibition downregulates mismatch repair genes, which may prime BRAF^V600E^ CRC with anti-PD-1 antibodies ^60^. A phase II trial of nivolumab (anti-PD-1 antibody) (NCT04017650) ^61^ or panitumumab (anti-PD-1 antibody) (NCT03668431) combination with MAPK inhibitor treatment demonstrated an improved response in MSS BRAF^V600E^ CRC patients ^62^. Our study utilized multiple CRC PDX models as key model systems on the basis of their advantage of close mimicry of patient tumor features but missing information about the immune microenvironment. The impact of the combination of BET and MAPK inhibition (e.g., BRAFi/EGFRi) on immune microenvironment reprogramming could be a novel area of future exploration.

Additionally, we observed dedifferentiation of cancer cells upon BRAF + EGFR or BET + BRAF + EGFR combinatorial inhibition in BRAF^V600E^ CRC PDX tumors. This finding suggests tumors experience cellular plasticity under therapeutic stress, which may result in a treatment-resistant tumor population and residual disease. However, it is critical to note that snMultiome sequencing was performed on tumor tissue samples collected after 10 days of treatment, which coincided with the onset of tumor regression under combination therapy to avoid the potential limitation of viable cell counts at the treatment endpoint. Intriguingly, we observed continuous tumor regression from day 10 to day 21, which indicates that BETi treatment may resolves the MAPKi-induced plasticity observed in CRC after an extended period of treatment. This study is the first to show that epigenomic reprogramming occurs during cell dedifferentiation alongside targeted therapy in BRAF^V600E^ CRC. Epigenetic reprogramming of dedifferentiated cell populations upon standard targeted therapy could be a novel area of exploration.

Overall, in this study, we identified abnormal chromatin dynamics associated with BRAF mutation in CRC and proposed a promising combination of epigenetic therapy with a standard targeted regimen for patients harboring BRAF^V600E^-mutated CRC. We are performing a further clinical assessment of ZEN-3694 + encorafenib + cetuximab in a phase I trial in treatment-refractory BRAF^V600E^ metastatic CRC (ClinicalTrial.gov identifier: NCT06102902).

## Methods

### CRISPR drug modifier screening

A Cas9-expressing BRAF^V600E^-mutated CRC cell line (RKO) was transfected with the sgRNA pool library containing selected genes associated with epigenomic regulation ^28,29^. The epigenomic library contains 519 epigenomic genes, 56 non-essential genes (negative control), and 73 core essential genes (positive control) (Supplementary Table #). Following puromycin selection for 72 hours (1 µg/ml), the initial cell population was harvested and cultured under the desired screening conditions for 5 population doubling times in a replicate of two (*n* = 2). The transfected cells were treated with either IC20 of vemurafenib (1 µM) or vemurafenib + cetuximab (0.1 mg/ml) and DMSO control for an additional 5 doubling times. sgRNA-barcoded sequences integrated into the genomic DNA of each cell population were amplified and subjected to high-throughput sequencing. DrugZ software ^63^ was used to obtain the gene-level normalized Z-score, which indicates the synthetic lethality of each gene under vemurafenib or vemurafenib + cetuximab treatment compared to DMSO control.

### Ribonucleoprotein-mediated CRISPR KO experiment

CRISPR KO of the BRD2 gene was conducted in two BRAF^V600E^ CRC cell lines, RKO and B1003 (PDX-derived cell line). We used Cas9-gRNA ribonucleoprotein methods, and ribonucleoproteins were prepared and delivered into cells using lipofection following the manufacturer’s protocol from Synthego. After 5 days of incubation, cell pools were collected and seeded onto 96-well plates to generate single KO clones. The sgRNAs (CRISPR evolution sgRNA EZ Kit) and Cas9 2NLS nuclease were purchased from Synthego, and Lipofectamine^TM^ CRISPRMAX Cas9 transfection reagent (CMAX00001) and Opti-MEM^TM^ I reduced serum medium (31985062) were purchased from Thermo Fisher Scientific. The sequences used for the sgRNA design were as follows: BRD2 sgRNA: cuuguuguaaauguaacagu, KIF11 sgRNA (essential positive control): ggaacuucacaacuuauugg, AAVS(T1) sgRNA (non-essential negative control): guccccuccaccccacagug.

### PDXs

Primary human-tumor PDX models were established as previously described ^64^. Tumor specimens were obtained from patients with metastatic CRC at The University of Texas MD Anderson Cancer Center (Houston, TX); all patients provided informed written consent for specimens to be used for research purposes, including implantation in PDXs. Samples were obtained with the approval of the institutional review board (IRB). All *in vivo* experiments and procedures were conducted with the approval of the Institutional Animal Care and Use Committee, and *in vivo* experiments involving PDXs were conducted following NIH NCI recommendation protocols in SOP50102: PDX Implantation, Expansion, and Cryopreservation (subcutaneous). Xenografts were established in 6- to 8-week-old female NOD/SCID mice purchased from Jackson Laboratory (Bar Harbor, ME, USA). Once established, PDXs were expanded in 6-week-old female athymic nude mice, purchased from Envigo Rms, Inc. (Indianapolis, IN, USA), subcutaneously. After tumors were established, with a median tumor volume exceeding 200 mm^3^, 3-10 mice/arm were treated via oral gavage with either vehicle control (EA006 solution [10% polyethylene glycol-300, 2.5% Tween 80 in water] or 0.5% Tween 80/0.5% CMC in water), or drugs, as indicated in the figure legends. Based on our prior experience with drug combination studies and datasets generated, sample size analysis reveals that the number of PDX models used in this study (with at least 3 tumors per treatment) is recommended to minimize random effect variance. ZEN-3694, iBET-151, and binimetinib were obtained via the CTEP agreement (U54 NCI), while encorafenib was purchased from Med Chem Express. Tumor sizes and mouse weights were measured twice a week. After 21 days, treatment was discontinued, and tumors from 3 mice per arm were excised (2-4 hours post-treatment), segmented, and immediately flash-frozen in liquid nitrogen (for protein, RNA, and DNA analysis) or 10% buffered formalin solution (for immunohistochemical staining).

### PDXOs

The BRAF^V600E^ CRC PDX model (B8302) with acquired resistance to encorafenib (BRAFi) + cetuximab (EGFRi) treatment was selected to establish the PDXO model. Fresh PDX tumor tissue was digested with tumor digestion media, which contains collagenase II (Thermo Fisher; 17101015), Dispase II (Thermo Fisher; 17105041), and FBS and then incubated, filtered, washed, and resuspended with phenol-red free Matrigel (Corning; 356231) before being seeded in a 24-well plate. After the Matrigel had fully solidified, organoid growth media was added to each well. After PDXO was established, organoids were seeded into 96-well black-wall plates (150 organoids/10 μl per well) in triplicate. After 3-5 days, when organoids started to grow, DMSO, encorafenib (0.5 μM) + cetuximab (0.1 mg/ml), ZEN-3694 (3 μM), or encorafenib + cetuximab + ZEN-3694 were added in fresh organoid growth media; cells were treated for 7 days. The CellTiter-Glo 3D Cell Viability Assay (Promega; G9683) was applied, and luminescence was measured as instructed. PDXOs seeded in 24-well plates (500 organoids/30 μl per well) were exposed to treatment in the same way and imaged using IncuCyte (Sartorius), per the manufacturer’s protocol. Organoid growth media was prepared with advanced Dulbecco’s modified Eagle medium/F-12 medium (Gibco; 12634010), GlutaMAX (Thermo Fisher; 35050061), HEPES (Thermo Fisher;15630080), Primocin (InvivoGen; ant-pm-2), A83-01 (Fisher Scientific; 29-391-0), recombinant human epidermal growth factor (Fisher Scientific; 236-EG-200), recombinant human Noggin (Fisher Scientific; 6057-NG-025), recombinant human fibroblast growth factor 10 (Fisher Scientific; 345-FG-025), gastrin I (Fisher Scientific; 30-061), N-acetylcysteine (Sigma-Aldrich; A9165-5G), nicotinamide (Sigma-Aldrich; N0636-100G), B-27 supplement (Fisher Scientific; 17504044), Y-27632 (Fisher Scientific; 12-541-0), and SB202190 (Selleckchem; S1077). L Wnt-3A and Rspondrin 1 cells were obtained from ATCC and R&D Systems, respectively, and Wnt3A- and Rspondrin1-conditioned media were collected following the vendor’s instructions.

### RNA-seq analysis

The STAR (v2.7.2b)^65^ RNASeq alignment tool was used for mapping raw BAM/FASTQ files to the human reference genome (GRCh38). Before alignment, Biobambam (v0.0.191) was used for indexing and marking duplicates ^66^. The Genecode (v22)^67^ gene annotation (.gtf) file was used in the data processing pipeline, and HTSeq (v0.11.0)^68^ was used to extract raw read counts from STAR-aligned files. Human tumor–derived sequence reads from mouse-derived reads in PDX model samples were classified using Xenome (v1.0.1-r) ^69^. Extracted human reads were processed as described.

### Differential gene expression analysis

Wald tests were performed in DESeq2 (v1.36.0)^70^ which performs normalization on samples and identifies differentially expressed genes between treatment comparisons. Non-expressed genes were filtered before normalization. RUVseq (v1.30.0)^71^ was used to correct hidden batch effects and remove unwanted variation from samples. Genes with an absolute log2 fold-change > 1.5 and adjusted *P* < 0.05 were identified as differentially expressed genes. A gene set enrichment analysis was conducted using genes that had been pre-ranked with shrunken log2 fold-change values as an input for fgsea (v1.22.0) ^72^ package in R. The Pheatmap package in R was used to plot variance-stabilizing transformation–normalized gene expression data. A principal component analysis and MDS plots between replicates of the same PDX model or treatment conditions were generated using ggplot2 ^73^ to examine non-technical variation and other latent factors.

### ChIP-seq analysis

BRD2 and BRD4 ChIP assays were performed at the MD Anderson Profiling Core, as previously described ^76^ with some modifications. Crosslinked tissue samples were sheared using optimized conditions for colon cells, with BRD2 (A302-583A) and BRD4 (A700-04) antibodies. Input and ChIP DNA libraries were prepared using an NEB Ultra II DNA library prep kit, following the manufacturer’s protocol, and subjected to next-generation sequencing to obtain ∼20 million 50-bp reads per sample. Human tumor–derived sequence reads were classified from mouse-derived reads in PDX model samples using Xenome (v1.0.1-r)^69^. Extracted human reads were processed using a snakemake-based ChIP-seq pipeline (https://zenodo.org/record/819971). Bowtie (v1.1.2)^77^ was used for the alignment of uniquely mapped reads to the human reference genome (GRCh37), and SAMBLASTER ^78^ was used for duplicate read removal. Uniquely mapped reads were downsampled per condition to 15 million, sorted, and indexed using samtools (v1.5) ^79^. deepTools v21.4.0 ^80^ was used to generate bigwig files to visualize data on the IGV genome browser by scaling the bam files to reads per kilobase per million. A model-based analysis of the ChIP-seq (MACS) (v1.4.2) ^81^ peak calling algorithm was used to define peaks. Enrichment of the peak was calculated over the “input” background genome data via a *P* value < 1e-5. Enhancers were assigned to genes using GREAT ^82^. Differentially bound peaks between each treatment group were identified using DiffBind ^83,84^. The Consensus Molecular Database Pathway analysis tool ^85^, Enricher ^86^, clusterProfiler ^87,88^ and GREAT were used for the pathway analysis. Using normalized bigwig files, a compressed score matrix for each PDX model was generated using the deeptools (v3.1.3) multibigwigsummary module.

### Super-enhancer-regulated TF analysis

Enhancers were identified using the Rank Ordering of Super-Enhancers algorithm ^89,90^. Enhancer constituents were stitched and classified as SEs in the rank-ordered H3K27ac ChIP-seq signal, normalized to input samples. CRCmapper ^38,39^ was used as previously described to computationally identify autoregulated SE-driven expressed TFs. MEME ^91^ software was used for the motif enrichment analysis of SE-associated TFs.

### MPAS analysis

The transcriptional level of MAPK signaling activity was measured using MAPK-specific 10 downstream gene targets (PHLDA1, SPRY2, SPRY4, DUSP4, DUSP6, CCND1, EPHA2, EPHA4, ETV4, and ETV5) that were identified and whose scores were calculated as previously described ^31^. The count matrix from H3K27Ac ChIPseq data was used and normalized for the 10 gene targets to calculate the MAPK signaling activity score. The Welch and Brown-Forsythe ANOVA test was performed using GraphPad Prism.

### Single-nucleus Multiome (RNA + ATAC) sequencing analysis

We performed single-nucleus Multiome (snRNA + snATAC) seq on a BRAF^V600E^ CRC PDX model (C5002) treated with control (Vehicle), BETi (ZEN-3694), BRAFi + anti-EGFR, or the triplet combination (two tumors per group) for 10 days. Single-cell suspensions from BRAF^V600E^ CRC PDX models were collected and processed using the 10X Genomics Multiome kit, as indicated in the manufacturer’s protocol (Rev A), for generating RNA-seq and ATAC-seq libraries. We targeted 6,000 cells per sample, and raw sequencing data were pre-processed and aligned to a chimeric hg38 and mm10 reference genome using cellranger-ARC v2.0.0 (10X Genomics), as previously described ^92^. We annotated cellular barcodes on the basis of the fraction of reads aligned to either the hg38 or mm10 genome. We set a cutoff of more than 70% of reads aligned to the human genome, more than 95% to the mouse genome for species annotation in RNA-seq, and 95% to either the human or mouse genome for species annotation in ATAC-seq. The cells that did not pass the cutoff were considered mixed human-and-mouse doublet combinations. The application of a joint RNA and ATAC filtering step resulted in 4,813 high-quality human cells in both modalities.

Batch effects were evaluated by k-BET ^93^ and corrected by Harmony ^93^ and Seurat v4 ^94^. Raw unique molecular identifier counts were log-transformed and used for principal component analysis. Seurat was applied to identify highly variable genes for unsupervised cell clustering, and UMAP was used for visualization ^95^. The weighted Nearest Neighbor (WNN) ^96^ analysis was used for integrated multimodal RNA + ATAC data analysis. Singac^97^ was used to analyze chromatin datasets, and chromVAR^98^ was used to identify enriched motifs. A cell cluster analysis using specific markers showed that several cells were characterized by different intestinal cell compartments (e.g., stem-like, transit-amplifying, and colonocyte-like cells). The differentiation states of each cluster of cell were computationally predicted using trajectory inference methods ^99^, including trajectory analysis (Monocle3 ^100^).

### Histology procedures and imaging

PDX tumor tissues were fixed in 10% neutral-buffered formalin for 48 hours, paraffin-embedded, sectioned 3- to 4-μm thick with a microtome, and placed on positively charged glass slides. After deparaffinization, hematoxylin-and-eosin (H&E) staining and immunohistochemical (IHC) staining were completed using a Leica Bond RX autostainer with a Leica detection kit. Brightfield whole-slide imaging was performed at 20x magnification with an Aperio AT2 scanner, and the images were analyzed by tuned Leica algorithms that were specific to each marker. Figures were generated using Imagescope v12.4.3.5008. The phosphoErk1/2 antibody (Cell Signaling Technology; 4370) was used for IHC staining.

### Statistical analyses

Statistical calculations were performed using GraphPad Prism version 8.0.0. Additional details are provided in the figure legends.

### Patient and Public Involvement

Patients and/or the public were not involved in the design, or conduct, or reporting, or dissemination plans of this research.

## Data availability

Raw and processed sequencing data generated in this study are available through Gene Expression Omnibus (GEO) at GSE280266 and GSE280280. The previously published CRC tumor ChIP-seq and HCT116 H3K27ac Hi-ChIP data were obtained from GSE88945. Additional raw or processed data used in this study are available upon request from the corresponding author.

## Author contributions

HML conceptualized and designed the study, performed the experiments, analyzed data, and wrote the manuscript. ZZ, CWW, SC, and SS performed computational analysis. AS and PMK performed PDX experiments. SN and VK performed experiments, provided technical support, and reviewed manuscript. AKS, CB, MP, EA, JYA, OV, VKM, JPS, AKJ, NWF, AA, DM, and AKS provided technical support and reviewed the manuscript. FMB provided resources and funding acquisition. KR and SK supervised and conceptualized the study, designed the experiments, and reviewed the manuscript. All the authors edited the manuscript.

## Ethics Statement

This study does not involve human participants. This study involves animal subjects only. Tumor specimens were obtained from patients with metastatic CRC at The University of Texas MD Anderson Cancer Center (Houston, TX); all patients provided informed written consent for specimens to be used for research purposes, including implantation in PDXs (LAB100982). Samples were obtained with the approval of the institutional review board (IRB). All in vivo experiments and procedures were conducted with the approval of the Institutional Animal Care and Use Committee, and in vivo experiments involving PDXs were conducted following NIH NCI recommendation protocols in SOP50102.

## Acknowledgments

This work was supported by the Cancer Center Support Grant – Gastrointestinal Program (P30CA016672), Gastrointestinal Research Foundation, SPORE program (P50CA221707), and the Department of Defense (DOD) Grant (CA230846). We also acknowledge CPRIT RP200390, ACS research scholar award (RSG-19-187-01-DMC), and internal grants for MDACC Epigenomics Therapy Initiative (METI) to K.R.. Next-generation sequencing (NGS) services were performed by The University of Texas MD Anderson Cancer Center Advanced Technology Genomics Core, generously supported by CA016672 and NIH 1S10OD024977-01. OEV was supported by the CPRIT Training Program (RP210028). AKJ was supported by institutional funding for MDACC Epigenomics Profiling Core. We thank Melanie Woods for technical support and useful discussions.

