## Supplemental Figure for "Reprogramming of Cellular Plasticity via ETS and MYC Core-regulatory Circuits During Response to MAPK Inhibition in BRAF-mutant Colorectal Cancer"

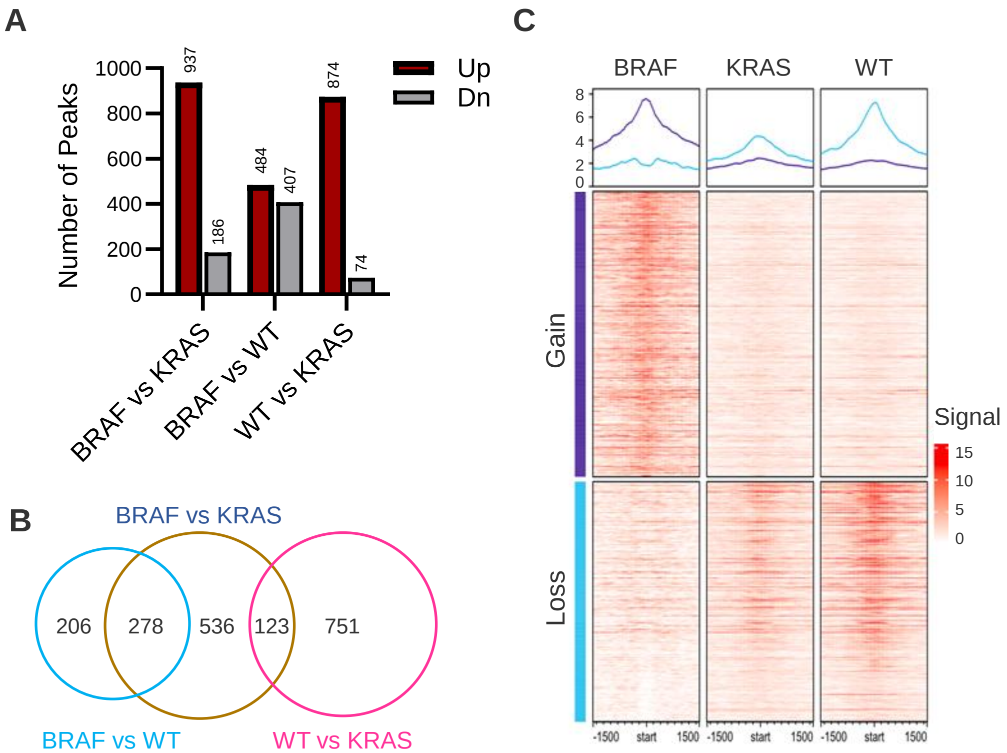


**Supplemental Figure 1. Differential H3K27ac enrichment among BRAF$^{V600E}$, KRAS$^{mut}$, and RAS/RAF wild-type CRC patient tumor samples.** (**A**) The number of differentially bound H3K27ac ChIP-seq peaks in three comparisons, such as BRAF$^{V600E}$-mutant CRC (BRAF; $n$ = 4) vs KRAS-mutant CRC (KRAS; $n$ = 7), BRAF-mutant CRC vs RAS/RAF wild-type CRC (WT; $n$ = 10), and WT CRC vs KRAS-mutant CRC, either upregulated (UP) or downregulated (DN) (adjusted $P < 0.01$). (**B**) Venn diagram of the number of significantly upregulated peaks in each comparison (BRAF vs KRAS, BRAF vs WT, WT vs KRAS; adjusted $P < 0.01$). (**C**) Profile enrichment of three groups of samples (BRAF, KRAS, and WT) around differentially bound peak sites from the BRAF vs WT comparison (gain: peaks enriched in BRAF-mutant samples, loss: peaks enriched in WT samples).

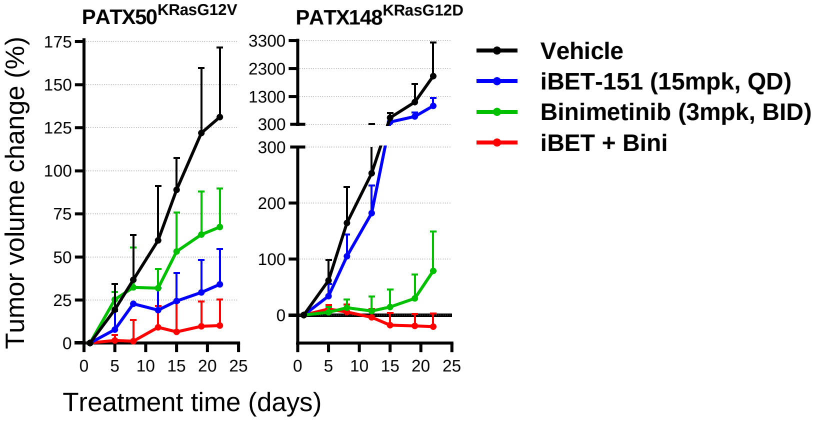


**Supplemental Figure 2. Expended iBET-151 + binimetinib combination screen in KRAS$^{mut}$ PDAC PDX models.** Tumor volume change relative to baseline was measured for 21 days for vehicle (control)-, binimetinib-, iBET-151-, and iBET-151 + binimetinib–treated PDAC PDX models (PATX148 and PATX50).

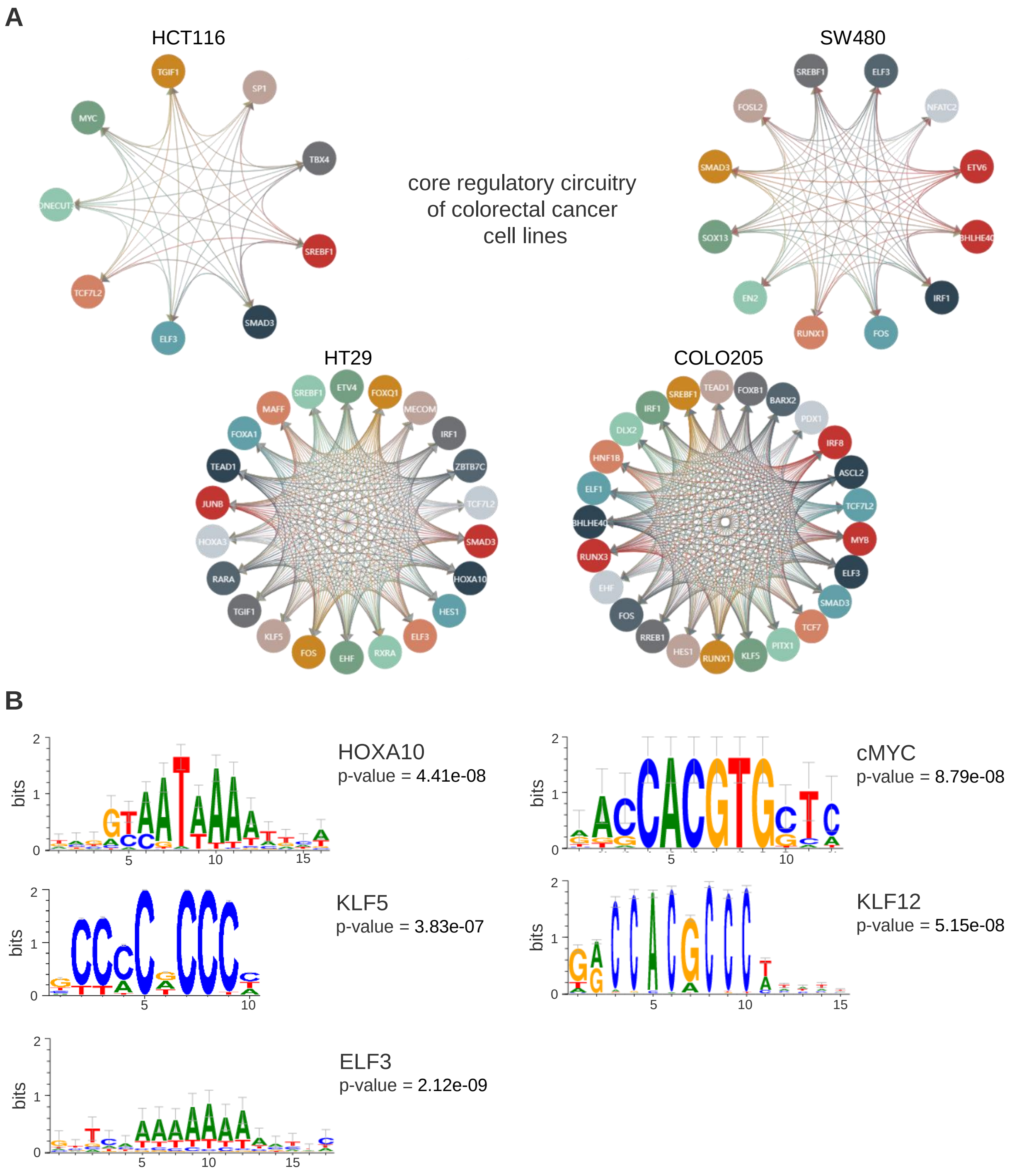


**Supplemental Figure 3. Core regulatory circuitries of CRC cell lines and PDX models.** (**A**) Core regulatory circuitries, identified from four CRC cell lines (HCT116, SW480, HT29, and COLO205) using a database of core transcriptional regulatory circuitries (dbCoRC; http://dbcorc.cam-su.org/) that is based on publicly available H3K27ac ChIP-seq datasets. Circles represent each TF connected with interactive loops. (**B**) Motif analysis of key auto-regulatory TFs (HOXA10, MYC, KLF5, KLF12, and ELF3) using H3K27Ac ChIP-seq data.

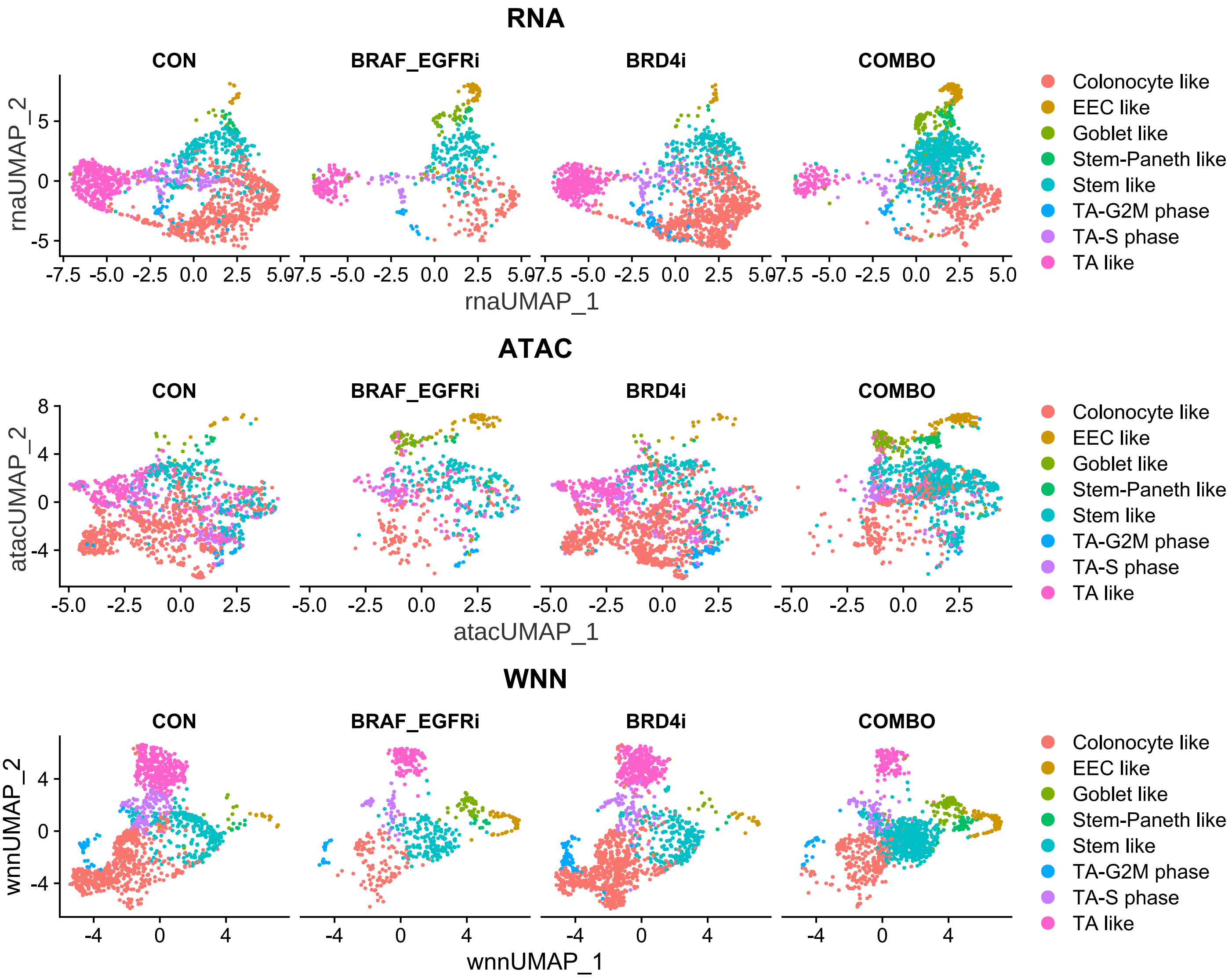


**Supplemental Figure 4. Differential abundance of each cell population by treatment.** The RNA, ATAC, and WNN combined UMAP projections of cell clusters, colored by different cell types in each treatment group, including control (CON), encorafenib + cetuximab (EC), ZEN-3694 (ZEN), and encorafenib + cetuximab + ZEN-3694 (ECZ).

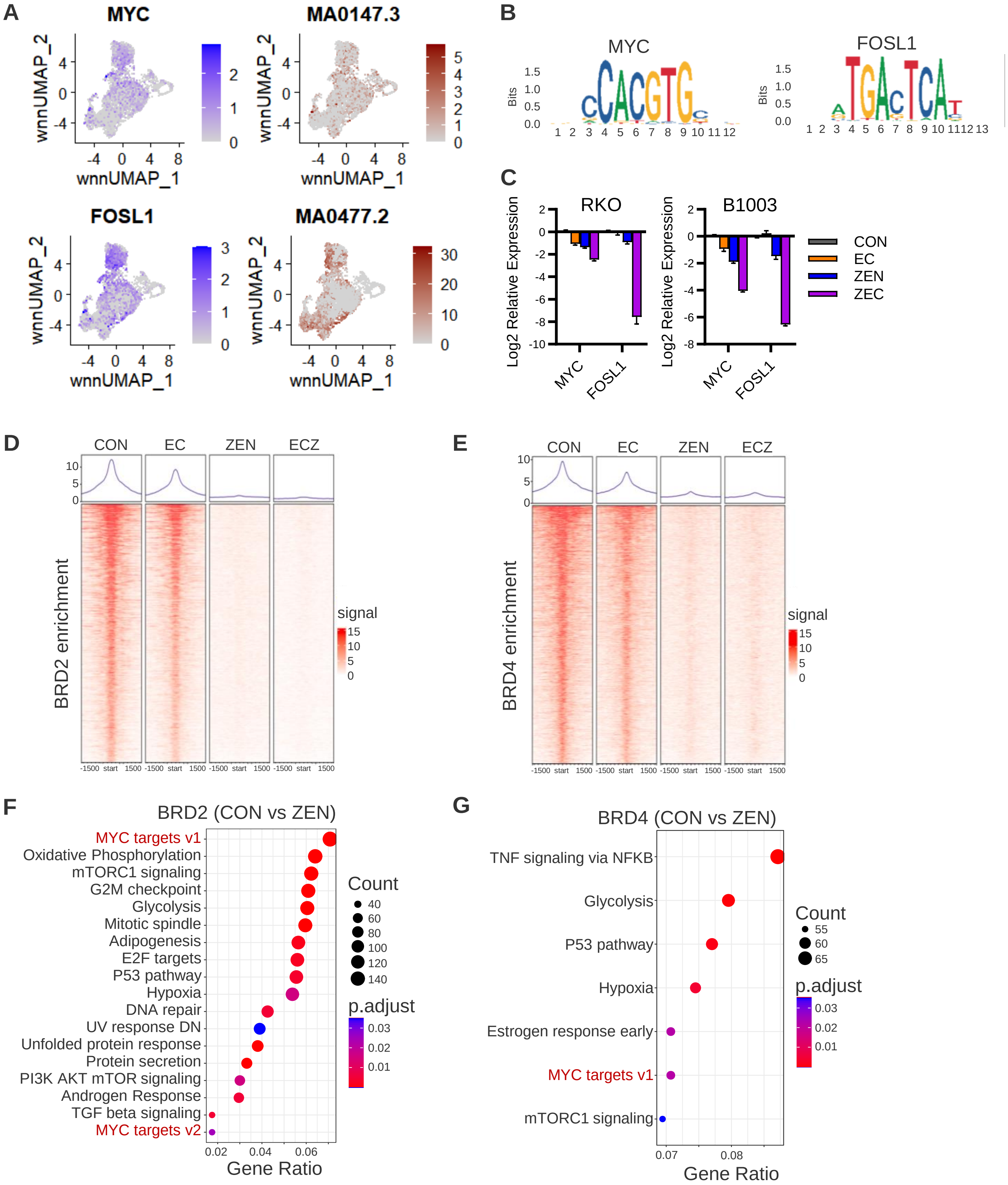
A
MYC
MA0147.3
FOSL1
MA0477.2
wnnUMAP_1
wnnUMAP_2
B
MYC
FOSL1
Bits
C
RKO
B1003
Log2 Relative Expression
MYC
FOSL1
CON
EC
ZEN
ZEC
D
CON
EC
ZEN
ECZ
BRD2 enrichment
signal
-1500 start 1500
E
BRD4 enrichment
F
BRD2 (CON vs ZEN)
MYC targets v1
Oxidative Phosphorylation
mTORC1 signaling
G2M checkpoint
Glycolysis
Mitotic spindle
Adipogenesis
E2F targets
P53 pathway
Hypoxia
DNA repair
UV response DN
Unfolded protein response
Protein secretion
PI3K AKT mTOR signaling
Androgen Response
TGF beta signaling
MYC targets v2
Count
p.adjust
Gene Ratio
G
BRD4 (CON vs ZEN)
TNF signaling via NFKB
Glycolysis
P53 pathway
Hypoxia
Estrogen response early
MYC targets v1
mTORC1 signaling
Count
p.adjust
Gene Ratio

**Supplemental Figure 5. BET inhibition blocked both BRD2 and BRD4 binding at a global level and around Myc target genes.** (**A**) WNN combined UMAP projection of the expression level of MYC and FOSL1 and enrichment of its associated motifs, MA0147.3 and MA0477.2, respectively, from C5002 PDX snMultiome-seq. (**B**). Motif logo of MYC (MA0147.3) and FOSL1 (MA0477.2). (**C**) Relative expression of MYC and FOSL1 in $BRAF^{V600E}$-mutated CRC cell lines (RKO and B1003) normalized to GAPDH, housekeeping gene, in different conditions, control (CON), encorafenib + cetuximab (EC), ZEN-3694 (ZEN), and triplet combination (ECZ) after 72 hours. (**D**) BRD2-normalized mean read count distribution of randomly selected 1k sites in each condition. (**E**) BRD4-normalized mean read count distribution of randomly selected 1k sites in each condition. (**F**) Enrichment of Hallmark gene sets with differentially enriched BRD2 peaks in control compared to ZEN-3694-treated samples (FDR < 0.05). Myc target gene sets are highlighted in red. (**G**) Enrichment of Hallmark gene sets with differentially enriched BRD4 peaks in control compared to ZEN-3694-treated samples (FDR < 0.05). Myc target gene sets are highlighted in red.
